# BRN2 preserves the melanocyte lineage following ionizing radiation

**DOI:** 10.64898/2026.10.18.759918

**Authors:** Jeremy H Raymond, Madeleine Le Coz, Zackie Aktary, Marie Pouteaux, Michael Hamm, Evelyne Sage, Vincent Favaudon, Valérie Petit, Alfonso Bellacosa, Colin R Goding, Lionel Larue

## Abstract

Adult stem cells maintain tissue homeostasis, but mechanisms preserving their integrity under stress remain incompletely understood. Here, we identify BRN2 (POU3F2), previously implicated in neuronal development, reprogramming of fibroblasts to neurons and in melanoma proliferation, migration and DNA-damage responses, as a key guardian of melanocyte stem cells. Remarkably, we reveal that while Brn2-deficient mice maintain their McSC compartment and normal pigmentation for up to 95 weeks, in contrast, rapid, dose-dependent, permanent hair depigmentation associated with the disappearance of detectable McSCs and differentiated melanocytes occurs in Brn2 KO mice exposed to ionizing radiation but not in WT mice. Importantly, our results indicate that in human follicular melanocytes BRN2 activity in melanocyte stem cells (McSCs) declines with age and that Brn2 promotes melanocyte stem cell regeneration by restraining stress-induced melanocytic differentiation. Our findings establish a previously unrecognized *in vivo* requirement for BRN2 in preserving the integrity of the melanocyte lineage following ionizing radiation, revealing a stress-dependent function distinct from its largely dispensable role in steady-state McSC maintenance.

## Introduction

Stem cells play a critical role in tissue homeostasis, enabling the regeneration of damaged tissues, replacement of cells with high turnover, and long-term self-renewal. Failure of these essential functions contributes to aging. Over the past decade, significant advances have been made in understanding the genes that drive stem cell activation and define stem cell niches (Khlusov et al. 2025; Zhang and Chen 2024). However, far less attention has been paid to how stem cell integrity is maintained in response to stress. Identifying factors that safeguard stem cells is critical, as inadequate protection can lead to stem cell depletion, tissue dysfunction, and ultimately loss of organ function.

Melanocyte stem cells (McSCs) provide an excellent model to study these processes. McSCs reside in the bulge of the hair follicle mini-organ and are responsible for generating melanocytes that maintain hair pigmentation (Hsu and Fuchs 2021; Nishimura 2011). During embryonic development, pigment-producing melanocytes are derived from melanoblasts originating from the neural crest, which contribute to the first round of hair pigmentation (Larue et al. 2013; Vandamme and Berx 2019). Postnatally, McSCs regenerate melanocytes required for hair pigmentation during successive hair cycles. At the start of anagen phase, McSCs proliferate and give rise to transit-amplifying cells (TACs), which migrate to the mature hair follicle bulb and differentiate into melanocytes, transferring pigment to adjacent keratinocytes (Bertrand et al. 2020; Nishimura et al. 2002). In mice, the first anagen occurs around one week after birth, the second around five weeks, and the third around 14 weeks.

The failure of this pigmentation system during aging is visible as hair graying and eventual whitening. Both genetic and environmental factors contribute to this process. For instance, loss of BCL2 impairs McSC survival, promoting hair greying (Nishimura et al. 2002), and deletion of β-catenin in adult McSCs similarly results in hair whitening (Le Coz et al. 2021; Rabbani et al. 2011; Takeo et al. 2016). Environmental stressors, including UV or ionizing radiation, accelerate hair graying through the generation of reactive oxygen species (ROS) that damage and eventually kill hair follicle melanocytes (Inomata et al. 2009; Ueno et al. 2014; Mohri et al. 2025). The microphthalmia-associated transcription factor (MITF) is central to the establishment and maintenance of the melanocyte lineage, coordinating cell survival, differentiation, proliferation, senescence, metabolism, and DNA repair (Goding and Arnheiter 2019). MITF activity is tightly regulated by activators such as SOX10, PAX3, β-catenin/LEF1, and CREB, as well as repressors including DEC1, ALX3, GLI2, FOXD3, and ATF4. One enigmatic transcription factor that can both upregulate and suppress MITF, depending on cellular context, is BRN2 (Goding and Arnheiter 2019; Goodall et al. 2008; Wellbrock et al. 2008).

BRN2, a member of the POU family of transcription factors, is expressed in the brain, including hypothalamic cells, is required for correct brain development (Andersen and Rosenfeld 2001; Fane et al. 2019; Nakai et al. 1995; Schonemann et al. 1995), and is critically required for reprogramming of fibroblasts into neuronal cells (Vierbuchen et al. 2010). In the melanocyte lineage, BRN2 expression is not detected in melanoblasts *in vivo* during the embryonic stages examined, although BRN2 is expressed in cultured melanoblasts (Goodall et al. 2004a; Berlin et al. 2012a; Goodall et al. 2004b). Although expressed in a several cancers such as prostate, glioblastoma, and small cell lung cancer (Schreiber et al. 1992; Bishop et al. 2017; Ishii et al. 2013; Suvà et al. 2014), its function has been most investigated in melanoma where BRN2 contributes to melanoma initiation, invasion, and proliferation and functions as a non-canonical tumor suppressor (Goodall et al. 2008; Fane et al. 2019; Berlin et al. 2012a; Eisen et al. 1995; Hamm et al. 2021; Zeng et al. 2018). In cultured melanoma cells, BRN2 can either activate or repress MITF expression depending on cellular context (Goodall et al. 2008; Wellbrock et al. 2008). More recently, a BRN2–MYC transcriptional axis has been identified as a regulator of melanoma phenotypic plasticity, with BRN2-high states associated with invasion and therapy resistance (Zhang et al. 2025). Notably, BRN2-high states were also detected in normal melanocytes. However, whether BRN2 regulates MITF-associated melanocytic differentiation in normal adult hair follicles *in vivo* remains unclear, especially since to date BRN2 expression has not been detected in the melanocyte lineage in vivo. BRN2 has also been implicated in the DNA-damage response, including regulation of GADD45 following UV irradiation (Lefort et al. 2001) and interacts with Ku70/Ku80 in melanoma cells (Herbert et al. 2019). However, how Brn2 influences the response of normal McSCs to ionizing radiation remains unknown. Notably, while UV and ionizing radiation induce overlapping damage responses, UV predominantly produces photoproducts whereas ionizing radiation can induce DNA double-strand breaks; the consequences for melanocyte-lineage fate therefore cannot be assumed to be identical.

Despite its established functions in melanoma and DNA-damage responses, whether Brn2 is expressed in the melanocyte lineage *in vivo* or whether it controls melanocyte differentiation or preserves adult McSCs following ionizing radiation remains unknown. Here, we investigate Brn2 expression in the melanocyte lineage *in vivo* and analyzed its function using melanocyte-specific Brn2 knockout mice. Our results reveal a critical physiological role for BRN2 as a guardian of McSCs following ionising irradiation, but not under steady state conditions. Our results are consistent with BRN2 restraining stress-driven melanocyte differentiation to allow regeneration of McSCs and prevent permanent hair depigmentation. Importantly, our results distinguish the physiological role of BRN2 in normal adult melanocytes *in vivo* from its previously described functions in cultured melanoma cells.

## Results

### Brn2 activity is enriched in melanocyte stem cells and declines with aging

Melanocyte stem cells (McSCs) maintain hair follicle pigmentation throughout life, and depletion of this population leads to hair greying and loss of follicular pigmentation. To identify candidate transcriptional regulators of McSC identity, we performed a comprehensive computational reanalysis of the previously published mouse single-cell RNA-sequencing dataset from Berns and colleagues (Berns et al. 2024). This reanalysis, rather than generation of the underlying single-cell dataset, is part of the present study and provided the rationale for focusing on Brn2.

Unsupervised clustering allowed for clear distinctions between different cell populations within the mouse hair follicle (Fig. 1A). By evaluating key melanocytic and stem cell markers, this clustering resolved a melanocytic compartment comprising the three established hair follicle melanocyte lineage populations: McSCs in the bulge, transient amplifying cells (TACs) in the intermediate region, and differentiated melanocytes in the bulb (Fig. 1B,C, Supplemental Fig. S1A,B).

**Figure 1.**
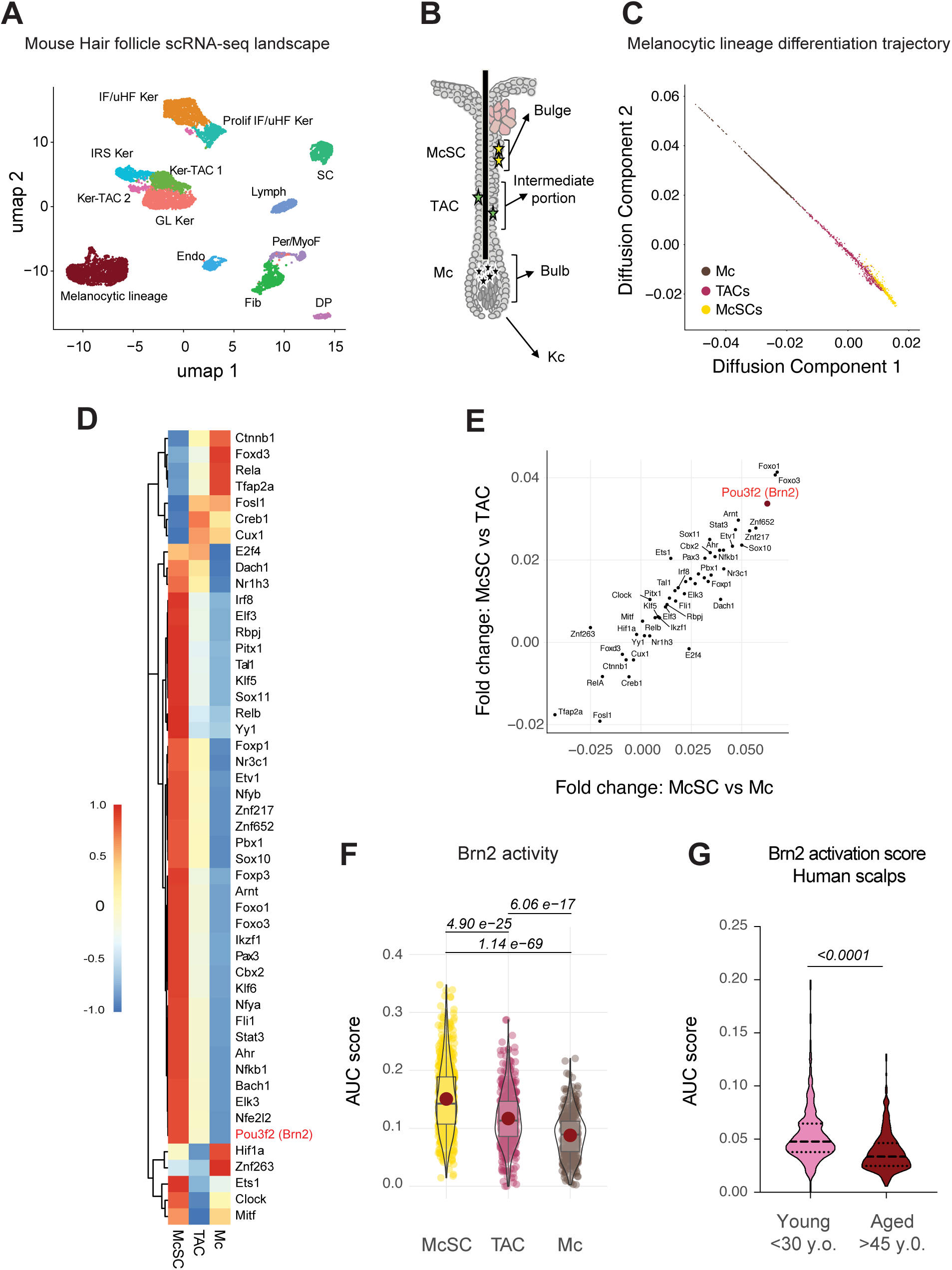
Transcription factor activity in the melanocytic lineage of mouse hair follicles. **A.** UMAP representation of the previously published mouse skin scRNA-seq (Berns et al.) showing major cell types: Endo: Endothelial Cells; Lymph: Lymphatic Vessel Cells; Per/MyoF: Pericytes/Myofibroblasts; Fib: Fibroblasts; DP: Dermal Papilla; SC: Schwann Cells; IFE/uHF Ker: Interfollicular/upper Hair Follicle Keratinocytes; Prolif IFE/uHF Ker: Proliferative Interfollicular/upper Hair Follicle Keratinocytes; GL Ker: Germinative Layer Keratinocytes; Ker-TAC: Keratinocyte Transit Amplifying Cells; IRS Ker: Inner Root Sheath Keratinocytes. **B.** Schematic of the hair follicle depicting the localization of Melanocyte Stem Cells (McSCs), Transit Amplifying Cells (TAC); Melanocytes (Mc), and Keratinocytes (Kc) **C.** Diffusion map of the mouse hair follicle melanocytic lineage **D.** Heatmap of average transcription factor activity for McSCs, Mel-TACs, and Mel. **E.** Correlation of fold changes in transcription factor activity scores between McSC *vs.* TAC and McSC *vs*. Mel; POU3F2 (Brn2) is highlighted. **F.** Brn2 activity score across melanocytic lineage cell types. Statistical analysis: ANOVA with Holm correction. **G.** Brn2 activity in human scalp melanocytic cells according to patient age category. Statistical analysis: ANOVA with Tukey post-test. This analysis shows an age association and does not establish a causal role for Brn2 in physiological hair greying.

To determine which transcription factors were active in each of these populations, we calculated transcription factor activity scores for individual cells using curated target gene sets associated with the melanocytic lineage (Fig. 1D). Comparison of these scores between McSCs, TACs and Mc allowed for the identification of transcription factors preferentially active in McSCs. Among these transcription factors was Brn2 (Pou3f2), which displayed one of the strongest activity enrichments in McSCs compared to other melanocyte lineage cells (Fig. 1D-F). Notably, analysis of an independent human scalp dataset revealed that Brn2 activity was lower in melanocytic cells from older individuals (Fig. 1G). This age-related association prompted us to investigate the functional role of Brn2. However, the absence of spontaneous greying in non-irradiated Brn2-deficient mice indicates that our findings do not establish a causal role for Brn2 in physiological hair aging in mice.

### Brn2 is not expressed in the melanocyte lineage during embryonic development

Previous studies have shown that Brn2 while BRN2 is expressed in cultured melanoblasts or cultured neural crest (Cook et al. 2003; Goodall et al. 2004a), its expression is absent in melanoblasts at embryonic day 11.5 (E11.5) (Goodall et al. 2004a). Whether it is expressed in melanoblasts at other times in development is not known. To examine whether Brn2 is expressed at later stages of melanocyte development, we used a mouse reporter line (Tyr::Cre; Z/EG^f/+^) in which melanoblasts express GFP (Colombo et al., 2012). GFP-positive melanoblasts were isolated from embryos between E14.5 and E18.5, and Brn2 mRNA levels were assessed by RT-qPCR. Brn2 transcripts were not detected at any of these stages (Fig. 2A). Pten served as a positive control for RNA quality and amplification, while cultured melanoblasts (Melb-a) served as a positive control for BRN2 expression.

**Figure 2.**
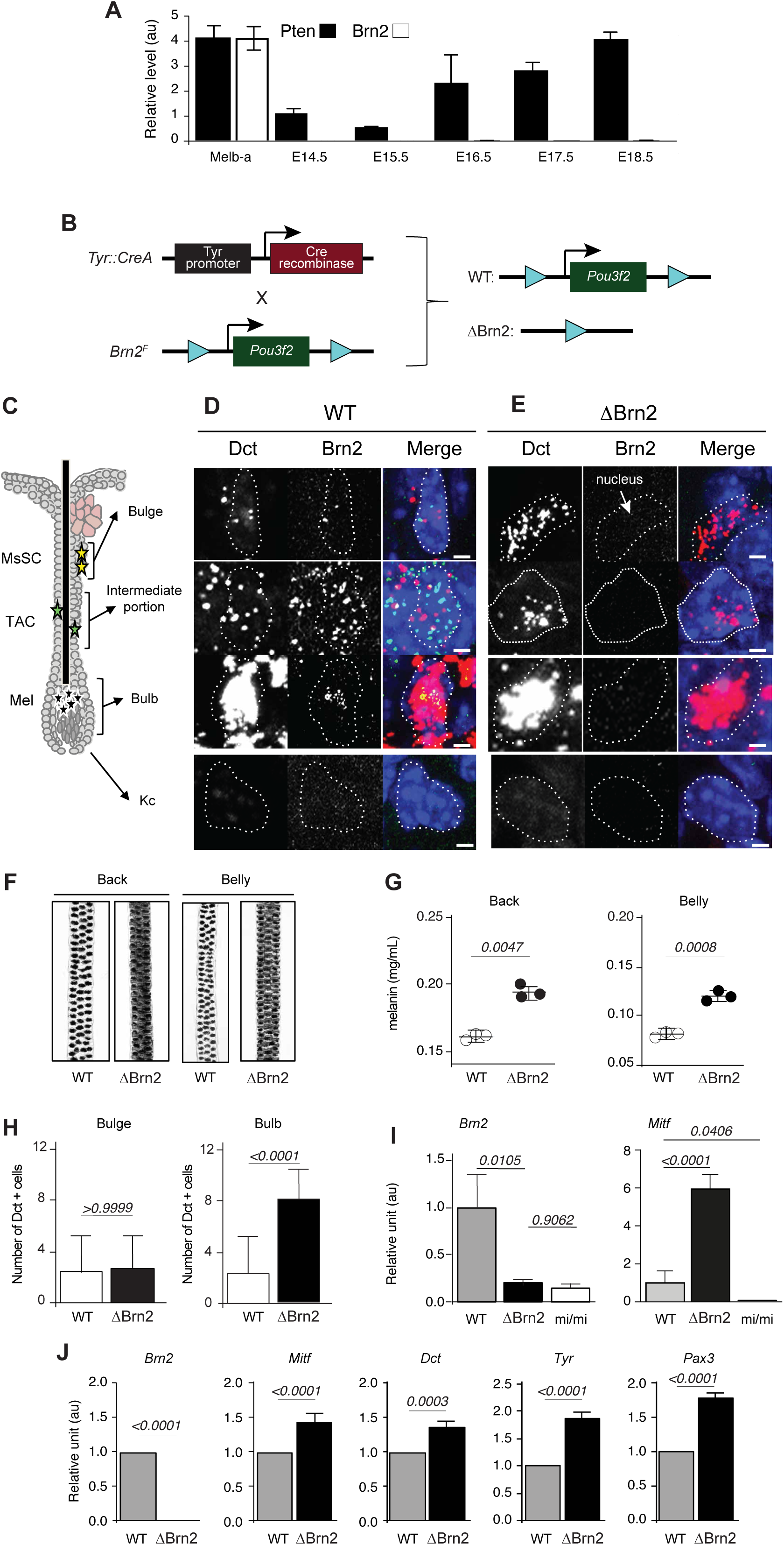
Effects of Brn2 loss in the melanocytic lineage. **A.** RT-qPCR analysis of Brn2 and Pten mRNA levels in GFP+ mouse melanoblasts isolated from embryonic skin during development. Three independent experiments were performed at each stage, with samples analyzed in triplicate. Melb-a cells were used as a cultured melanoblast control. Hprt was used for normalization. **B.** Schematic representation of the mouse model used to generate melanocytic lineage-specific Brn2 deletion (ΔBrn2 = Tyr::Cre; Brn2 ^F/F^, and WT = Brn2^F/F^). **C.** Diagram of an anagen hair follicle showing the anatomical location of melanocytic lineage cells: melanocyte stem cells (McSC) in the bulge, melanocyte transit-amplifying cells (Mel-TAC) in the intermediate region of the follicle, and differentiated melanocytes (Mel) in the bulb. Keratinocytes (Kc) and the sebaceous gland are also indicated. **D. E.** RNAscope detection of Dct and Brn2 mRNA in hair follicle sections from wild-type (WT; Tyr::Cre; Brn2^+/+^) (d) and Brn2-deficient (ΔBrn2; Tyr::Cre; Brn2^F/F^) mice (e). Confocal microscopy images show Dct (red), Brn2 (green), and nuclei (DAPI, blue). Dots correspond to individual mRNA molecules. Dct marks melanocytic lineage cells. The dotted white line delineates a nucleus. Scale bar = 1 μm. **F.** Representative images of hairs from the back and belly of adult WT and ΔBrn2 mice. **G.** Quantification of melanin content in hairs collected from back and belly skin of WT and ΔBrn2 mice. Data are from three independent biological experiments. Statistical significance assessed using an unpaired *t*-test with Welch correction. **H.** Quantification of Dct+ melanocytic lineage cells in the bulge and bulb regions of WT and ΔBrn2 hair follicles. Statistical significance was assessed using an unpaired t-test using data obtained from at least three independent embryos and at least three follicles per sample. **I.** RT-qPCR analysis of Brn2 and Mitf mRNA levels in back skin from postnatal day 4 (P4) WT, ΔBrn2, and melanocyte-deficient Mitf ^vga9/vga9^ (mi/mi) mice. Data are from four independent biological experiments. Statistical significance was assessed using an unpaired *t*-test with Welch correction. **J.** RT-qPCR quantification of melanocytic lineage genes (Mitf, Dct, Tyr, Pax3) in melanocyte cultures derived from WT and ΔBrn2 mice. Gapdh was used as internal control. Error bars represent s.d. Data are from four independent biological experiments. Statistical significance was assessed using an unpaired *t*-test.

These results indicate that, unlike melanoblasts or neural crest in culture, Brn2 is not expressed, or is expressed at levels below detection, in melanoblasts *in vivo* during embryonic development. This suggests that Brn2 may function predominantly postnatally in the melanocyte lineage.

### Brn2 loss in the melanocytic lineage promotes melanocyte accumulation in the hair follicle bulb

Since constitutive *Brn2 (Pou3f2)* knockout mice die within the first 10 days after birth due to severe hypothalamic defects (Nakai et al. 1995), we generated mice with a melanocyte-specific deletion of *Pou3f2*, the gene encoding BRN2 (*Tyr::Cre; Brn2^F/F*), to determine the functional role of BRN2 in the melanocyte lineage (Fig. 2B, Supplemental Fig. S2A,B). RNAscope analysis confirmed efficient loss of Brn2 expression in melanocytic cells throughout the hair follicle, including the bulge, intermediate region, and bulb (Fig. 2C-E).

Although the overall coat color of mutant mice appeared similar to controls and independent of the number of hair follicle cycles (Supplemental Fig. S2C-F), histological analysis revealed a clear increase in melanin deposition within hair follicles lacking Brn2 (Fig. 2F,G). This hyperpigmentation was associated with a marked increase in the number of Dct-positive cells in the bulb, corresponding to differentiated melanocytes, whereas the number of Dct-positive cells in the bulge (McSCs) remained unchanged (Fig. 2H). While surprising, this observation is consistent with human data that showed that human follicles are darker with age in patients not subject to hair greying (Commo et al. 2012; Itou et al. 2019). Importantly, this increase in differentiated melanocytes was not accompanied by depletion of the stem-cell compartment: the number of Dct-positive cells in the bulge remained unchanged (Fig. 2H), and the coat color of ΔBrn2 mice remained indistinguishable from that of controls throughout life, up to 95 weeks of age, irrespective of the number of hair cycles undergone (Supplemental Fig. S2C-F). Because each hair cycle requires renewed McSC activation and production of differentiated melanocytes, progressive exhaustion of the stem-cell pool would be expected to result in a cycle-dependent loss of pigmentation. The absence of such a phenotype indicates that the increased differentiated output associated with Brn2 loss does not compromise long-term maintenance of the McSC compartment under steady-state conditions. Brn2 is therefore dispensable for maintenance of the melanocyte lineage in the unchallenged follicle. Moreover, repeated depilation-induced hair cycles imposed no detectable pigmentary cost in either genotype, indicating that increased proliferative demand alone is insufficient to deplete the McSC pool within the lifespan of a mouse. Thus, replicative challenge and genotoxic stress have distinct consequences for this lineage, and the permanent depigmentation observed following irradiation cannot be explained simply by cumulative proliferative history.

In agreement with this observation, transcriptomic analyses revealed an age-associated increase in the melanocytic gene expression signature in humans (Supplemental Fig. S2G). The specificity of Brn2 expression for the melanocyte lineage in the hair follicle was proven by analysing mouse skin lacking melanocytes (*Mitf^mi/mi^ mice*) by RT-qPCR (Fig. 2I). Moreover, RT-qPCR analysis of established melanocytes from ΔBrn2 mice revealed an increased expression of melanocytic genes upon Brn2 deletion (Fig. 2J). Together, these results suggest that Brn2 normally restrains differentiation of the melanocyte stem cell lineage.

### Brn2 deficiency alters DNA-repair-associated transcriptional programs in melanocytic cells in melanocytic cells

To investigate the molecular mechanisms underlying these phenotypes, we performed RNA-seq analysis of control and Brn2-deficient melanocytic cells. Analysis revealed that while the Brn2-low cells increased the level of expression of genes associated with matrix remodeling, their proliferation and DNA repair pathways were downregulated (Fig. 3A). Gene set enrichment analysis (GSEA) revealed a significant downregulation of DNA repair pathways in the absence of Brn2, largely driven by reduced expression of key genes involved in nucleotide excision repair (NER), including *Xpa, Xpc, and Ercc1*, and chromosome maintenance genes such as *Smc2* and *Smc4* (Fig. 3B,C), suggesting that Brn2 supports the expression of DNA-repair- and chromosome-maintenance-associated transcriptional programs in melanocytic cells. To test whether Brn2-deficient cells display increased sensitivity to genotoxic stress, we examined their transcriptional response following ionizing radiation (IR).

**Figure 3.**
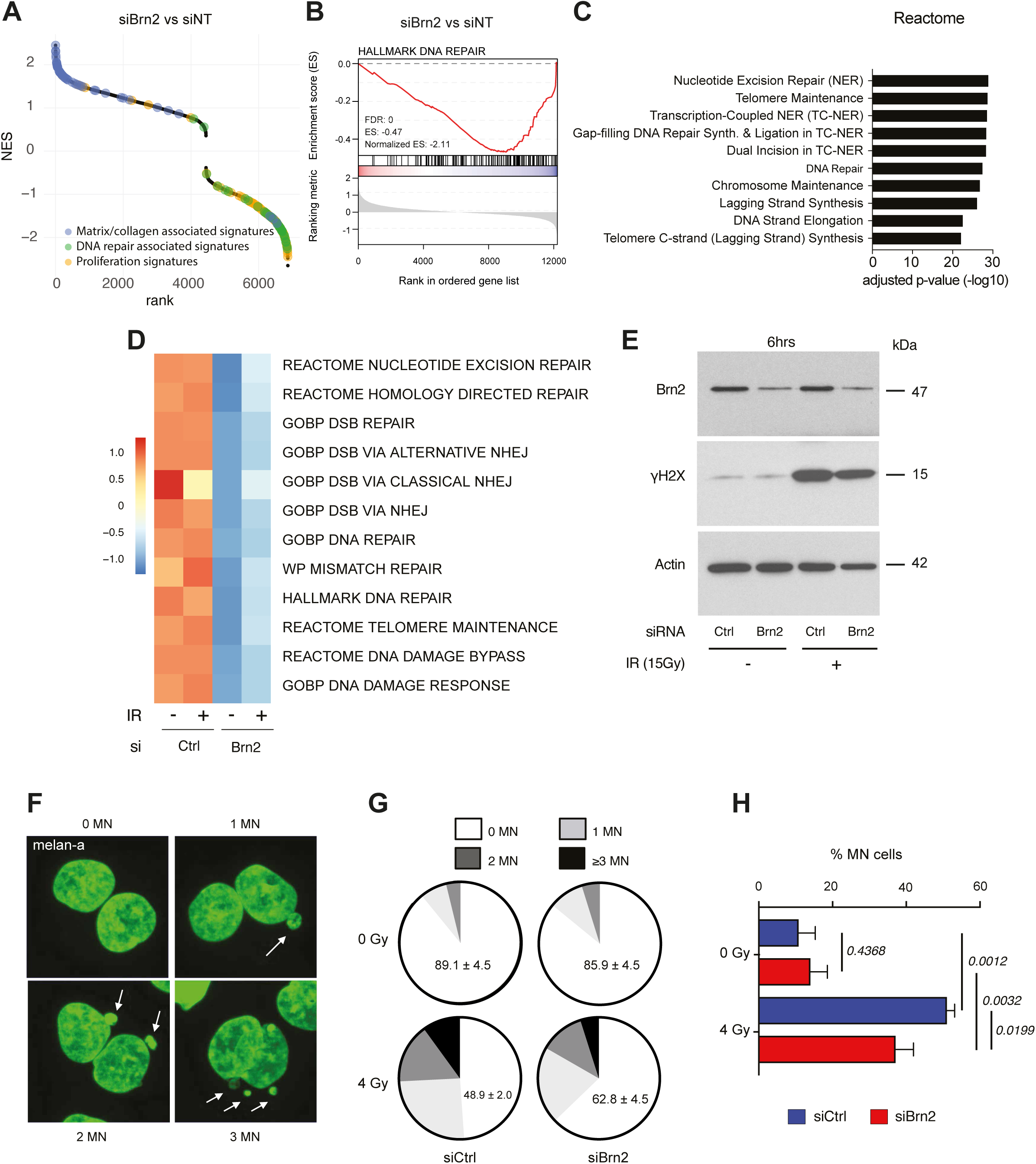
Brn2 regulates DNA repair in melanocytes. **A.** GSEA NES of all MsigDB-C2 gene sets from of siBrn2 versus siNT Melan-a, ranked by NES. Matrix/collagen-associated (blue), DNA repair-associated (green) and proliferation-associated (orange) gene sets are highlighted; positive NES denotes enrichment among genes upregulated after Brn2 knockdown. **B.** Gene Set Enrichment Analysis (GSEA) of DNA-repair signatures in melan-a cells transfected with control siRNA (siCtrl) or Brn2-targeting siRNA (siBrn2). **C.** Reactome enrichment analysis of leading-edge genes from the DNA-repair signature. **D.** Heatmap showing average expression levels of core nucleotide excision repair (NER) and double-strand break (DSB) response/repair genes in melan-a cells following Brn2 knockdown. **E.** Western blot analysis of Brn2 and γ-H2AX protein levels in control and Brn2-depleted melan-a cells after ionizing radiation. Actin was used as a loading control. This blot is interpreted as supportive evidence of altered DNA-damage-response signaling and not as a quantitative measurement of DNA-repair efficiency. **F.** Representative images of nuclei displaying different numbers of micronuclei following irradiation. **G.** Distribution of micronuclei categories in control and Brn2-depleted melan-a cells after irradiation. **H.** Percentage of cells containing micronuclei under control or Brn2 knockdown conditions following irradiation. Statistical significance was assessed using Chi-squared tests with PRISM software based on data from three independent biological experiments.

The loss of Brn2 induced a repression of genes involved in most of the repair pathways independently of IR (Fig. 3D). However, when looking at the core genes of these pathways – such as *Xpa*, *Xpc, Ercc1*, *Ercc2*, *Trp53bp1, or Xrcc4*, IR seems to potentialize the activity of Brn2 (Supplemental Fig. S3A). Consistent with these transcriptional alterations, the γH2AX immunoblot showed a lower irradiation-associated signal in Brn2-depleted cells (Fig. 3E). Because this single immunoblot does not by itself provide a quantitative measure of DNA repair efficiency, we now interpret it only as supportive evidence of altered DNA-damage-response signaling and base our conclusion primarily on the transcriptomic DNA-repair signatures. A blunted γH2AX response in cells with elevated MITF is, moreover, the expected outcome rather than an anomaly: upon DNA damage MITF is redirected into a complex with MRE11–RAD50–NBS1 and impairs MRN recruitment to damaged DNA (Binet et al. 2024). Because Brn2 represses Mitf^16^, Brn2 loss may therefore attenuate double-strand-break signalling indirectly, through MITF de-repression, in addition to reducing the transcription of nucleotide-excision-repair genes.

Despite the altered DNA-repair-associated transcriptional profile, Brn2-deficient cells exhibited significantly fewer micronuclei following irradiation (Fig. 3F-H; Supplemental Fig. S3B-D). Coupled with the decreased proliferative signature observed in melanocyte cells with decreased Brn2 levels (Fig. 3A), these data suggest that cells lacking Brn2 may exit the proliferative cycle rather than accumulate chromosomal instability. Because micronuclei can only be scored in cells that have completed a division, and fewer Brn2-depleted cells remain proliferative following irradiation (Fig. 4D,E), the reduced micronucleus frequency likely reflects, at least in part, the smaller cycling population rather than improved repair of DNA damage. Thus, this finding should not be interpreted as evidence that Brn2 loss enhances genome maintenance.

**Figure 4.**
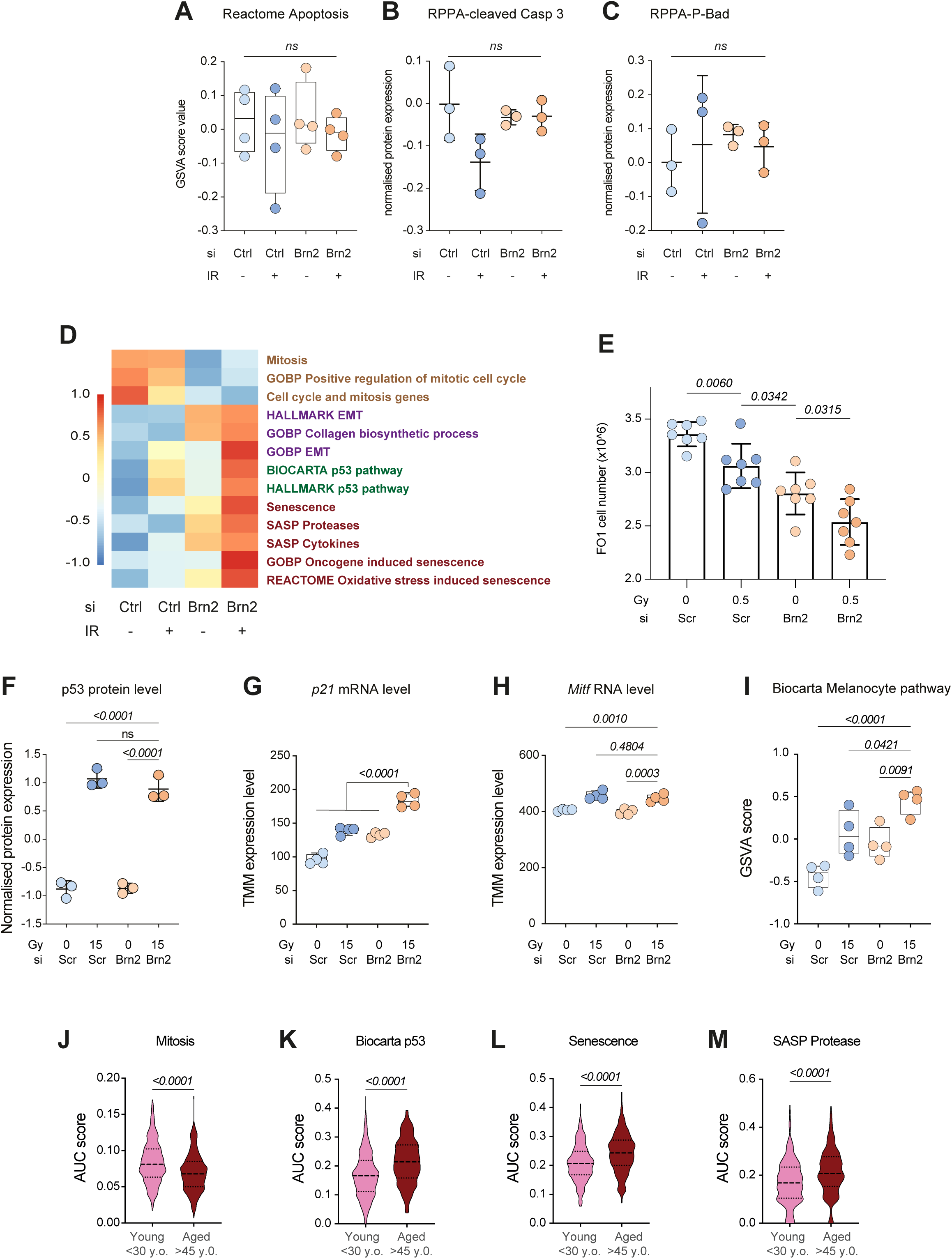
Brn2 modulates apoptosis-associated, mitotic, and stress-induced differentiation in melan-a cells. **A.** Gene Set Variation Analysis (GSVA) scores of the Reactome Apoptosis signature computed from bulk RNA-seq in melan-a cells following Brn2 knockdown and 15 Gy irradiation. **B,C.** Reverse Phase Protein Array (RPPA) quantification of cleaved caspase-3 (**B**) and phosphorylated Bad (**C**) in melan-a cells transfected with control or Brn2 siRNA and exposed to 15 Gy irradiation. **D.** Heatmap of mean GSVA scores for dysregulated signatures computed from bulk RNA-seq in melan-a cells following Brn2 knockdown and 0.5 Gy irradiation. **E.** Quantification of FO-1 cell numbers in cultures with the indicated combinations of Brn2 knockdown and 0.5 Gy irradiation. Statistical significance was assessed using an unpaired t-test based on data from at least six independent biological experiments. **F.** RPPA quantification of p53 protein levels in melan-a cells following Brn2 knockdown and 15 Gy irradiation, showing increased p53 after irradiation and a higher signal in Brn2-depleted irradiated cells. **G,H.** RNA-seq quantification of Cdkn1a (p21; G) and Mitf (H) expression levels in melan-a cells following Brn2 knockdown and irradiation. p21 induction is interpreted as evidence of p53-associated cell cycle control and is not considered a senescence-specific marker. **I.** Biocarta melanocyte pathway GSVA score in melan-a cells following Brn2 knockdown and 15 Gy irradiation. **J-M.** AUC scores for mitosis (J), p53 activation (K), senescence-associated (L) and SASP protease-associated (M) transcriptional signatures in melanocytic cells from human scalp according to patient age category. These are transcriptomic signatures and do not by themselves demonstrate cellular senescence. Statistical analysis: t-test. For panels **A-I** statistical comparisons were performed by Anova followed by a Tukey post-test comparing the siBrn2 + IR group with the other groups, unless otherwise indicated.

### Brn2-deficient cells display a p53-p21-associated stress-induced differentiation phenotype following genotoxic stress

To determine the fate of Brn2-deficient cells after irradiation, we first evaluated whether increased apoptosis accounted for the reduced micronuclei formation. However, analysis at the transcript and protein levels did not reveal significant differences the cell-death-related markers examined between control and Brn2-deficient cells (Fig. 4A-C). These measurements do not exclude transient or context-dependent apoptosis, particularly *in vivo*.

Instead, RNA-seq analysis revealed a marked reduction in mitotic gene signatures in Brn2-deficient cells (Fig. 4D), indicating decreased proliferative capacity. Furthermore, the number of cells in culture with Brn2 knockdown and irradiation was significantly lower than all other cultures (Fig. 4E).

Ionizing radiation induced robust expression of p53, to a similar extent in control and Brn2-depleted cells (Fig. 4F, Supplementary Table S1). In contrast to p53 itself, the p53 target and cyclin-dependent kinase inhibitor p21 (*CDKN1A*) was significantly upregulated in irradiated Brn2-depleted cells compared with all other conditions (Fig. 4G), consistent with enhanced cell-cycle arrest, although p21 induction alone is not specific for senescence. Importantly, irradiated cells lacking Brn2 expression displayed moderately increased levels of Mitf and a higher melanocyte gene expression signature (Fig. 4H,I). Interestingly, in human scalp melanocytic cells, mitotic gene-expression signatures decreased with age, whereas p53-, senescence- and SASP-associated transcriptional signatures increased (Fig. 4J-M). These transcriptomic associations do not demonstrate bona fide cellular senescence.

Together, these results show that Brn2-deficient melanocytic cells respond to genotoxic stress with reduced proliferation, activation of a p53–p21-associated program, and increased melanocytic differentiation. In the absence of complementary functional senescence assays, we do not conclude that these cells undergo bona fide senescence. Rather, we interpret this phenotype as a stress-induced, senescence-associated differentiation state, reminiscent of the recently described concept of “senodifferentiation” in stem cell systems(Mohri et al. 2025). The transcriptional response to irradiation was, moreover, coordinated and extended well beyond p21 induction. In irradiated Brn2-depleted cells, gene sets associated with senescence, SASP cytokines, SASP protease regulators, oncogene-induced and oxidative stress-induced senescence, as well as the BIOCARTA and HALLMARK p53 pathways, all reached their highest enrichment scores, whereas three independent mitotic gene sets reached their lowest (Fig. 4D). Importantly, these gene sets are largely non-overlapping, and neither SASP set contains cell-cycle inhibitors, indicating that this coordinated response cannot simply be attributed to *Cdkn1a* induction. Notably, p53 protein was induced by irradiation to a similar extent in control and Brn2-depleted cells (Fig. 4F), whereas downstream p53 transcriptional activity was more pronounced in the absence of Brn2. Thus, Brn2 loss appears to modify the transcriptional outcome of the p53 response rather than the magnitude of p53 induction itself. Moreover, Brn2 loss alone does not induce this programme but rather enhances the response of the melanocytic lineage once DNA damage has occurred.

These findings should, however, be interpreted in the context of the acute timepoints examined: RNA was collected one hour and protein six hours after irradiation. Since establishment of a bona fide senescent state typically requires a longer period, the absence at these early timepoints of protein-level features associated with established senescence, including changes in p16INK4a (CDKN2A), IL-6 and Rb phosphorylation in our RPPA dataset (Supplementary Table S1), is not unexpected. We therefore interpret these data as evidence for the early induction of a coordinated senescence-associated transcriptional programme rather than as evidence that the cells have already entered an established senescent state.

### Brn2 protects the melanocyte stem cell compartment from irradiation-induced depletion

Given the reduced expression of DNA-repair genes and the altered differentiation responses observed in Brn2-deficient cells, we hypothesized that Brn2 may be required to maintain the melanocyte stem cell pool under genotoxic stress. To test this hypothesis, control and Brn2-deficient mice were exposed to increasing doses of ionizing radiation (10, 15, and 20 Gy) once, at 7 weeks of age, and hair pigmentation was monitored over time.

Whereas control mice maintained largely normal pigmentation following irradiation, Brn2-deficient mice developed pronounced and dose-dependent depigmentation of the irradiated areas (Fig. 5A). Importantly, this depigmentation was permanent over the observation period, consistent with loss of regenerative melanocyte-lineage capacity in the irradiated area. Depigmentation was confined to the irradiated fields, appeared within the first weeks after exposure, and hairs regrowing in these areas remained unpigmented at every subsequent cycle examined (Fig. 5A).

**Figure 5.**
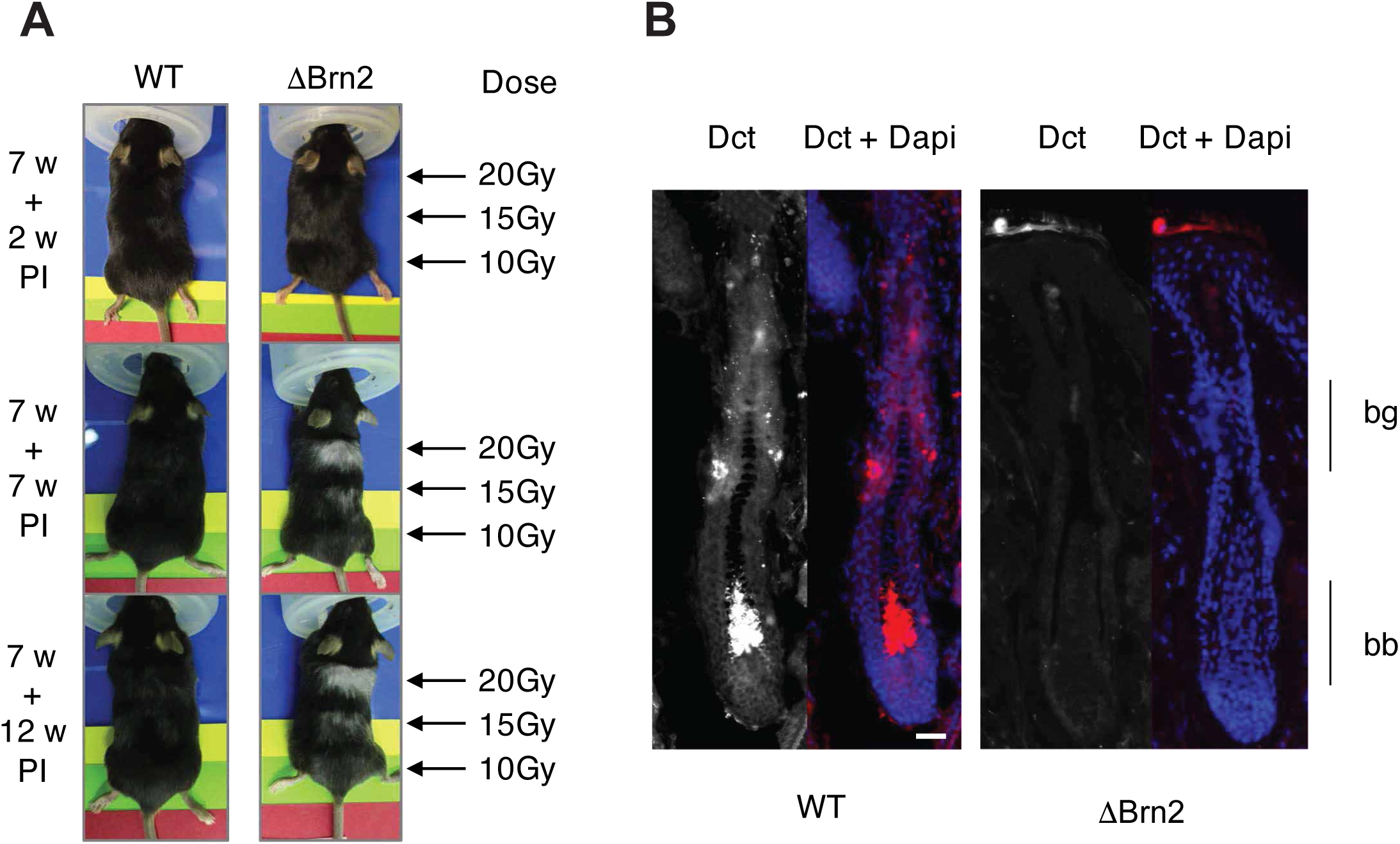
Brn2 deficiency sensitizes hair follicles to irradiation. **A.** Representative photographs of WT and ΔBrn2 mice at 2, 7, and 12 weeks following localized ionizing irradiation. Seven-week-old mice were irradiated on the dorsal skin using a linear accelerator (LINAC) at 10, 15 and 20 Gy. **B.** RNAscope staining for Dct in skin sections from WT and ΔBrn2 mice collected seven weeks after irradiation. Nuclei were counterstained with DAPI. Scale bar = 25 μm. bg = bulge; bb = bulb. The absence of Dct+ cells in irradiated DBrn2 follicles supports loss of the melanocyte lineage after genotoxic stress.

Consistent with this interpretation, RNAscope analysis revealed a complete absence of both McSCs and differentiated melanocytes in irradiated hair follicles of Brn2-deficient mice (Fig. 5B), whereas these populations were present in the irradiated hair follicles of control mice. These findings establish an *in vivo* requirement for Brn2 in preserving detectable melanocyte-lineage populations following ionizing radiation.

## Discussion

Hair follicle melanocyte stem cells (McSCs) maintain pigmentation throughout life, yet the mechanisms governing their response to genotoxic stress remain poorly understood. Here, we identify a physiological role for BRN2 (POU3F2) in the adult melanocyte lineage *in vivo*. BRN2 activity is enriched in McSCs and declines with age. Its deletion enhances melanocytic differentiation and reduces DNA-repair-associated transcriptional programs without compromising McSC maintenance under steady-state conditions. However, following ionizing radiation, Brn2-deficient mice exhibit McSC depletion and permanent hair depigmentation, associated with altered DNA-damage responses, reduced proliferation, p53-p21 activation, and enhanced differentiation.

Our findings reveal complementary functions of BRN2. First, BRN2 restrains melanocytic differentiation, as its deletion increases MITF-associated gene expression and differentiated melanocyte accumulation in hair follicles. Previous studies reported opposing effects of BRN2 on MITF expression in cultured melanoma cells, suggesting context-dependent regulation (Goodall et al. 2008; Wellbrock et al. 2008). Our findings extend these observations by demonstrating a physiological role for BRN2 in adult melanocyte differentiation *in vivo*.

Second, BRN2 supports DNA-repair-associated transcriptional programs, including nucleotide excision and double-strand break repair pathways. Unlike the non-transcriptional interaction of BRN2 with Ku70/Ku80 described in melanoma cells (Herbert et al. 2019), our findings implicate BRN2 in the transcriptional regulation of DNA-repair-associated genes. However, these changes do not establish direct regulation of DNA repair or progressive genomic instability. The increased differentiated output and preservation of McSCs under steady-state conditions are not contradictory. McSCs can compensate for increased differentiation through self-renewal and progenitor reversion (Sun et al. 2023). Thus, BRN2 may regulate the threshold for differentiation rather than being essential for homeostasis. Its requirement becomes apparent following irradiation, when McSC depletion prevents subsequent pigmentation regeneration. Similar stress-induced depletion has been reported in ATM-deficient McSCs (Inomata et al. 2009) and following DNA damage-associated senodifferentiation (Mohri et al. 2025). Our findings identify BRN2 as another determinant of McSC preservation under genotoxic stress.

Third, BRN2 is required for melanocyte-lineage preservation following ionizing radiation. Irradiated Brn2-deficient mice develop rapid, dose-dependent, and permanent hair depigmentation, accompanied by the disappearance of detectable McSCs and differentiated melanocytes. Following irradiation, Brn2-deficient cells exhibit altered γH2AX signaling, reduced mitotic activity, increased p21 expression, and enhanced differentiation. Interestingly, fewer micronuclei are detected, potentially reflecting reduced proliferation or cell-cycle exit rather than decreased DNA damage. These observations are compatible with a stress-associated differentiation response resembling senodifferentiation (Mohri et al. 2025).

Together, these findings support a model in which BRN2 preserves McSC integrity following genotoxic stress by modulating the balance between DNA-damage responses, melanocytic differentiation, and stem-cell maintenance. The observed senescence-like features and transcriptional changes in DNA-repair-associated pathways are consistent with a stress-associated differentiation response that may contribute to McSC depletion. Although additional mechanisms, including apoptosis, may also contribute, our results identify BRN2 as an important regulator of melanocyte stem-cell fate and regenerative capacity under genotoxic stress. Future studies will be required to elucidate the molecular mechanisms linking BRN2 to DNA-damage responses and to clarify the respective contributions of differentiation, senescence, and cell death to McSC depletion.

A comparable stress-dependent requirement has been described for another POU transcription factor, OCT1/POU2F1, in intestinal stem and progenitor cells. Oct1 deletion minimally affects steady-state homeostasis but compromises regeneration following tissue injury (Vázquez-Arreguín et al. 2019). Although OCT1 and BRN2 belong to distinct POU subclasses and respond to different stresses, these observations suggest that preservation of regenerative capacity under stress may represent a broader function of POU transcription factors.

Our findings also extend previous observations implicating BRN2 in melanoma DNA-damage responses, including Ku70/Ku80 interactions and protection against UVB- and chemotherapy-induced apoptosis (Herbert et al. 2019). Importantly, our study addresses normal adult melanocytes rather than transformed cells and identifies a genetically defined requirement for BRN2 following ionizing radiation *in vivo*. Unlike UV-induced photoproducts, ionizing radiation can generate DNA double-strand breaks, potentially triggering distinct cellular responses. Nevertheless, our results do not establish that differentiation replaces apoptosis, since transient cell death cannot be excluded from endpoint analyses.

Beyond McSC maintenance, these findings raise the possibility that BRN2 contributes to melanoma radioresistance through phenotype switching. A BRN2–MYC transcriptional axis was recently identified as a regulator of melanoma phenotypic plasticity, with BRN2-high states associated with invasion and therapy resistance (Zhang et al. 2025). Given BRN2’s involvement in melanoma plasticity and its association with genotoxic stress responses in normal melanocytes, we hypothesize that BRN2-expressing melanoma cells may exhibit enhanced resistance to ionizing radiation. Such a mechanism could contribute to melanoma radioresistance, although this hypothesis requires experimental validation.

Our study further identifies an association between BRN2 loss, p53-p21 activation, MITF-associated transcriptional activity, and altered DNA-repair-associated programs. However, the absence of p53 inhibition experiments prevents conclusions regarding its mechanistic requirement. Identifying direct BRN2 targets and determining its interactions with p53 and MITF will be essential to clarify the transcriptional mechanisms governing McSC responses to genotoxic stress.

Although BRN2 activity declines in melanocytic cells from older human scalp samples, our findings do not establish a causal role in physiological aging. Brn2-deficient mice maintain pigmentation during prolonged observation without irradiation, and their lifespan cannot reproduce decades of human hair aging. Thus, the demonstrated requirement for BRN2 concerns genotoxic stress rather than spontaneous age-associated greying.

In conclusion, our findings establish BRN2 as a regulator of melanocytic differentiation and stress-dependent preservation of adult McSCs in vivo. BRN2 restrains differentiation under physiological conditions and contributes to maintaining the melanocyte lineage following irradiation. Its loss favors stress-associated differentiation and permanent depigmentation, although the relative contributions of differentiation and cell death remain unresolved. These findings may have broader implications for stem-cell biology, as defective nucleotide excision repair causes premature aging syndromes, including xeroderma pigmentosum and Cockayne syndrome, while efficient genome maintenance has been associated with longevity (Firsanov et al. 2025; MacRae et al. 2015). Our results suggest that coordination between DNA-damage responses and cell-fate decisions may contribute to preserving stem-cell integrity under stress, although its relevance to physiological aging remains to be established.

## Materials and Methods

### Mouse hair follicle scRNA-seq analysis

Data from the analysis published by Berns and colleagues (Berns et al. 2024) were downloaded from GSE236918. Data were uploaded in R using Seurat (Hao et al. 2024) and cells with less than 2^10^ features and more than 20% reads mapping to mitochondrial genes were removed. Doublets were also identified and removed using scDblFinder (Germain et al. 2021). Data were integrated based on the first 100 dimensions using harmony. Marker genes were identified using seurat FindAllMarker function and a cell type was attributed to each cluster based on those identified genes. Melanocytic cells were identified based on the expression of Dct. Melanocytic cluster was subset and a diffusion map was created using destiny. Melanocytes were identified by the high expression of differentiated melanocyte genes such as *Oca2*, *Pmel*, and *Mc1r*. McSCs were identified by their highest expression of *Bcl2, Pax3,* or *Klf4*. TACs were identified as the cells in between Melanocytes and McSCs and are characterised by high proliferation scores. Transcription factor activity was computed using the AUCell function of the package escape (Borcherding et al. 2021). Transcription factor differential activity was generated using Limma, Data were represented using pheatmap, ggplot2 and ggstaplot (https://doi.org/10.1007/978-3-319-24277-4_9, https://doi.org/10.21105/joss.03167).

### Human scalp scRNA-seq analysis

Data from the analysis published by Ober-Reynolds and colleagues were downloaded from <u>GSE212450</u>. Data were uploaded in R using seurat and cells with less than 300 features and more than 10^4.5^ counts and 25% reads mapping to mitochondrial genes were removed. Data were integrated based on the first 100 dimensions using harmony. Melanocyte cluster was identified based on the expression of DCT. Transcription factor activity was computed using the AUCell function of the package escape.

### Generation of a transcription factor activity signatures in melanocytic cells

To generate an unbiased list of TF targets in melanocytic cells, TPM expression data from The Cancer Genome Atlas TCGA-SKCM were downloaded using TCGAbiolinks (Colaprico et al. 2016). The expression of each gene was correlated to all the other ones using pearson. Only genes with a correlation >0.4 with the «prey» gene were kept. Transcription factors (TFs) were subseted from this list and only the genes shown to be bound by the TFs in ChIP-seq analysis were kept. The list of ChIP-seq used was ChEA 2022. The list of transcription factors and their targets is provided in Supplementary Table S2.

### Transgenic mice

Tyr::Cre and Brn2 F/F were previously generated (Delmas et al. 2003; Jaegle et al. 2003). All transgenic mice were maintained on a C57BL6/J background. Animal care and use for this study were performed in accordance with the recommendations of the European Community (2010/63/UE) for the care and use of laboratory animals. Experimental procedures were specifically approved by the ethics committee of the Institut Curie CEEA-IC #118 (CEEA-IC 2016-001) in compliance with the international guidelines.

### Cell lines

Melanocyte cell lines were established in cell culture from the skin of 1.5 day old WT and ΔBrn2 littermate pups as previously described (Berlin et al. 2012b). WT, ΔBrn2 and Melan-a mouse melanocyte cells were all grown in F12 medium (Gibco, 21765-029) supplemented with 10% FBS (Gibco, 10270-106), antibiotics (100 U/mL penicillin and 100 μg/mL streptomycin; Gibco, 15140), and 200nM TPA (Sigma, P1585). FO-1 human melanoma cell lines were grown in RPMI 1640 media (GIBCO, 11875101), supplemented with 10% FBS and 1% antibiotics. All cell lines were maintained at 37°C in a humidified atmosphere containing 5% CO_2_.

### BRN2-CRISPR plasmid construction

CRISPR-Cas9-empty plasmids were a generous gift from Thomas Zillinger (University Hospital Bonn, Germany). CRISPR-BRN2-knockout and CRISPR-GFP-knockout plasmids were cloned by linearizing the CRISPR-Cas9-empty plasmid with *SwaI* (NEB, R0604) in NEB buffer 3.1 for 1hour at 25°C, followed by heat inactivation of the enzyme at 65°C for 20 minutes. A target-specific oligo was then annealed with the CRISPR Universal antisense oligo (each 100μM) and denaturated at 95°C for 5 min following a temperature gradient of 0°C/30s down to 8°C. The annealed oligos were filled up with 0.4μL of dNTP’s (10μM) and incubated at 37°C for 1hour followed by an inactivating step of 75°C for 10 min. 0.5μL of the filled in oligos were mixed with 1μL (15ng) of the purified linearized CRISPR-Cas9-empty plasmid, mixed with 1.5μL H_2_O and 2,5μL Gibson Assembly Master-mix (2x Gibson Master Mix [250μL]: 100μL 50% PEG 5000 [w/v], 50μL 1M Tris pH 7.5, 15μL H_2_O, 5μL 1M MgCl_2_, 10μL dNTP’s [10μM], 10μL 50mM NAD^+^ [NEB, B9007S], 5μL 1M DTT, 50μL Taq Ligase [40U/μL; NEB, M0208S], 6.25μL Phusion DNA Polymerase [2U/μL; NEB, MO530S] and 0.2μL T5 exonuclease [10U/μL; NEB, M0363S]) and incubated at 50°C for 1 hour. Two μL of this mix was then used to transform ultracompetent bacteria XL10 Gold (Stratagene, 200314) according to manufacturer’s instructions. The next day, clones were picked and grown over night on a shaker at 37°C in 5 mL LB medium with ampicillin (100 μg/mL). Plasmid DNA isolation was performed according to the manufacturer’s instructions of the Qiagen Quiaprep Spin Miniprep Kit (Qiagen, 27104). Plasmid preparations (1μg) were analyzed by agarose gel electrophoresis (AGE) after digestion with *Swa*I and *EcoR*V (NEB, 3195) in NEB buffer 3.1.

### CRISPR transfection and cell line generation

FO-1 cells were seeded on 10cm cell culture dishes and transfected at 60% confluency with 24μg of the respective CRISPR-Cas9 plasmid per plate using Lipofectamine 2000 (ThermoFisher, 11668019) according to the manufacturer’s protocol. After 48 hours, Blasticidine (4μg/mL) was added to the cell culture media and maintained for three weeks. Blasticidine-resistant cells were then trypsinized, washed and resuspended in PBS for single-cell sorting using the BD FACS ARIA III cell sorter. Single-cell clones were established and propagated in growth media containing Blasticidine (4μg/mL). Isolated clones were validated by western blot for presence/absence of BRN2 protein. Genomic DNA of selected clones was extracted using the AllPrep DNA/RNA Mini Kit (Qiagen, 80204) and subjected to DNA sequencing to determine the introduced mutation.

### RNAseq

For RNAseq experiment, melan-a mouse cells were transfected using lipofectamine 2000 (Invitrogen) with 100pmoL of siRNA targeting mouse BRN2 (Dharmacon M-020029-01-0005) or a Scrambled sequence. Cells were irradiated with GSR D1 irradiator Cs-137 at a dose of 15Gy. Transfections and irradiations were done 4 times on consecutive passages of the cells. RNA was extracted 1 hour after irradiation using the miRNAeasy kit (Qiagen, 217004), according to the manufacturer’s protocol. RNA integrity (RIN) was measured using an Agilent Bioanalyser 2100 and a RNAnano 6000 kit (5067-1511, Agilent Technologies). All RNA had a RIN of 10. RNA concentrations were measured using a NanoDrop (NanoDrop Technologies). RNA-sequencing libraries were prepared from 1μg total RNA using an Illumina TruSeq Stranded mRNA library preparation kit, which allows strand-specific sequencing. PolyA selection using magnetic beads was done to focus the sequencing on polyadenylated transcripts. After fragmentation, cDNA synthesis was done and the resulting fragments used for dA-tailing, followed by ligation with TruSeq indexed adapters. The fragments were amplified by PCR to generate the final barcoded cDNA libraries (12 amplifications cycles). The libraries were equimolarly pooled and subjected to qPCR quantification using the KAPA library quantification kit (Roche). Sequencing was carried out on a NovaSeq 6000 instrument (Illumina) based on a 2 x 100 cycle mode (paired-end reads, 100 bases) using an S1 flow cell. Reads were mapped to the mm10 mouse reference genome (gencode m13 version-GRCm38.p5) using STAR. STAR was also used to create the expression matrices. Reads were mapped to the mm10 mouse reference genome (gencode m13 version-GRCm38.p5) using STAR (Dobin et al. 2013). Batches were corrected with Combat (Leek et al. 2012). Differential gene-expression analysis was done following the DEseq2 pipeline (Love et al. 2014). GSEA was done using fgsea (https://doi.org/10.1101/060012) on signature at least 15 genes. Other signature scores were obtained by GSVA. The list of the signature used for the analyses is provided in Supplementary Table S3. Graphical representations were made with ggplot2 (https://doi.org/10.1007/978-3-319-24277-4) and pheatmap.

### RPPA

For RPPA experiment, melan-a mouse cells were transfected using lipofectamine 2000 (Invitrogen) with 100pmoL of siRNA targeting mouse BRN2 (Dharmacon M-020029-01-0005) or a Scrambled sequence. Cells were irradiated with GSR D1 irradiator Cs-137 at a dose of 15Gy. Transfections and irradiations were done 3 times on consecutive passages of the cells. Total proteins were extracted 6 hours after irradiation with the following lysis buffer : 1% Triton X-100, 50mM HEPES, pH 7.4, 150mM NaCl, 1.5mM MgCl2, 1mM EGTA, 100mM NaF, 10mM Na pyrophosphate, 1mM Na3VO4, 10% glycerol, containing freshly added Complete protease inhibitor (Roche, 11873580001) and PhosStop phosphatase inhibitor cocktail (Roche, 4906845001). Cell lysates were mixed with 4X sample buffer (40% glycerol, 8%SDS, 0.25M Tris-HCL Ph6.8, beta-mercaptoethanol), boiled for 5 minutes and store at -80°C until sample submission to the MD Anderson Cancer Center. Protein extracts were spotted onto nitrocellulose-coated slides arranged in a 4×12-pin-11×11 subarray that accommodate 1056 serially diluted samples plus replicate vertical controls on each slide. Proteins of interest were detected by probing with specific antibodies, amplifying the signal via a tyramide amplification system and visualized via DAB colorimetric reaction. The detection system was a GenPoint-based staining kit from Agilent. Digital images of slides were obtained by scanning on a Huron TissueScope scanner producing 16-bit TIFF files. Spot intensities from the TIFF files were determined via Array-Pro Analyzer software. Mean net intensities of spots from all 1056 samples were used for curve-fitting for each slide (RPPA Super Position and Concentration Evaluation, aka SPACE). Relative protein level for each sample was determined by converging its sample dilution series to one point (EC_50_) and interpolating in RPPASPACE. Expression was normalised sample loading differences by bidirectionally median centering values by antibody then by sample and then adjusted for Set-to-Set differences (batch effects) by normalizing identical control samples in this set with an invariant control sample set and applying the adjustment (differences in means × inverse of standard deviation ratio) to each corresponding data point.

### siRNA transfection and irradiation of cell lines

Melan-a and FO-1 cell lines were seeded on 6-well plates 24-hours prior to transfection. When at 70% confluence, cells were transfected using lipofectamine 2000 (Invitrogen) according to the manufacturer’s instructions with 1μg plasmids or 100pmoL of the scrambled or Brn2-specific siRNAs (Dharmacon). Six hours after transfection, the culture medium was changed.

Cells received 0.5 or 15 Gy γ-rays, depending of the cell type, using a ^137^Cs GSR D1 irradiator (Gamma-Service Medical GmbH) twenty-four hours after transfection. At specific time points post-irradiation, cells were rinsed twice with PBS, and protein or RNA was extracted as described below.

### siRNA sequences

siRNA targeting mouse and human *BRN2* were purchased from Dharmacon as a SMARTpool mix of 4 sequences. The sequences of the siRNA are as follows: for human and mouse *BRN2*: 5’-CGG AUC AAA ACU GGG AUU UA-3’, 5’-GCA AGG GCG CAA GCG GAA A-3’, 5’-AAG GAG GUG GUG AGA GUU U -3’, 5’-CGA CCU UUG CAG GCG AGU AA-3’. A random, scrambled sequence (5’-AAU UCU CCG AAC GUG UCA CGU-3’) was used as a control.

### Semi-quantitative Real-time PCR

Total RNA was extracted from cells using the miRNeasy kit from Qiagen (217004) according to the manufacturer’s protocol. Reverse transcription reactions were performed with 500 ng of total RNA and M-MLV reverse transcriptase (Invitrogen), according to the manufacturer’s instructions. The cDNA generated was then subjected to quantitative real-time PCR for the analysis of gene expression using ABI 7900HT. Oligonulceotides used for PCR are given in Supplementary Table S4.

### Western blot analysis

Whole cell lysates were prepared from melanocyte cell lines using ice-cold RIPA buffer supplemented with complete protease inhibitor cocktail and PhosStop phosphatase inhibitor cocktail (Roche, 11873580001). For western blotting, 25 μg total protein was separated on 15% denaturing acrylamide SDS-PAGE gels and the proteins transferred to nitrocellulose membranes. Membranes were blocked in 5% non-fat milk in Tris-buffered saline supplemented with 0.05% Tween-20 (TBST) and probed with primary antibodies overnight. The signal was detected using peroxidase-conjugated anti-mouse or anti-rabbit secondary antibodies (Jackson 111-035-003 and 111-035-045, respectively) and enhanced chemiluminescence (ECL; Pierce 34075). The primary antibodies used in this study are listed in Supplementary Table S5.

### Epilation of mouse hair

The overall coat color of the mice and color of individual hairs were evaluated as previously described at birth and during the life of adult mice. Hair depilation was performed on mice with a 1:1 mixture of beeswax (Sigma, 243221) and rosin gum (Sigma, R3755). Depilation was first done at seven weeks of age and every four weeks following the first epilation on the back of isofluoran-anesthethized mice. Pictures of the coat color were taken regularly after each epilation.

### Hair count, hair pictures and melanin content assay

Hair plucking was performed on the back and the belly of three mice of each genotype. Random hairs from each genotype were mounted between slides and coverslips in PBS containing 0.1% glycerol and observed under a light microscope. Pictures were taken with light microscope and assembled in Adobe Illustrator. The melanin content of plucked hair follicles was quantified as previously described (Berlin et al. 2012b).

### Irradiation of mice and isolation of skin biopsies

Seven week-old mice were irradiated on their backs using a linear accelerator (LINAC) device (Lansonneur et al. 2019) with 10, 15 or 20 Gy on different regions of the back. The coat color was documented with pictures over time. Hairs were plucked for observation through the life of the animal and skin biopsies were taken as previously described (Berlin et al. 2012b). Mice were euthanized and shaved prior obtaining skin biopsies. Skin was removed and either fixed in 4% PFA in PBS overnight or directly frozen in liquid nitrogen until RNA extraction. The fixed biopsies were included in OCT and frozen over liquid vapours. Cryosections of 10µm thickness were cut and mounted on SuperFrost Plus slides (Fisher Scientific 12-550-15) in the direction of the hair in order to obtain longitudinal sections of the hair follicles. These sections were used for RNAScope staining and immunohistofluorescence.

### RNAscope

Dct mRNA and Brn2 mRNA were detected in skin cryosections of WT and ΔBrn2 mice using RNAscope immunostaining technology following the manufacturer’s recommendations (https://acdbio.com/technical-support/user-manuals) Images were taken using a CLSM SP5 Leica confocal microscope and assembled using ImageJ and Adobe Illustrator.

### Immunofluorescence

Immunostaining of skin sample sections was done using samples fixed in 4% PFA prior to inclusion. All washing steps were done with 0.2% Tween 20 in PBS. Slides carrying skin cryosections were washed and antigene unmasking in citrate buffer was performed while boiling for 20min. The slides were washed again before blocking in PBS + 0.2% Tween 20 and 2% BSA for 1h. The slides were then incubated O/N in a humidified chamber at 4°C with primary antibodies against Dct (Santa Cruz, sc-10451) diluted in PBS + 0.2% Tween 20. The next day, the slides were washed and incubated for 1h at RT in the dark with the secondary antibodies Alexa Fluor 488 (Invitrogen, A21206). The sections were again washed before being incubated with 0.5 µg/µL DAPI to visualize the nucleus. Slides were washed and mounted in Vectashield mounting medium (Vector Laboratories) before imaging by epifluorescence or confocal microscopy. The images were assembled using ImageJ and Adobe Illustrator.

### Micronucleus assay

FO-1 cells were seeded at 1×10^5^ cells per well on glass coverslips in 6 well plates. FO-1 cell lines were transfected with Brn2 or Scrambled siRNAs 24 hours after seeding using Lipofectamine 2000. 24 hours after transfection, the cell lines were irradiated with 0Gy or 4Gy γ-rays from a ^137^Cs irradiator. For the FO-1-CRISPR-GFP and FO-1-CRISPR-BRN2 cells, irradiation was done 48hours after seeding. Immediately after irradiation, the cells were treated with 3μg/mL Cytochalasin B (diluted in fresh growth medium), which was maintained for 30 hours. Glass coverslips were then rinsed twice in PBS and fixed in 4% PFA in PBS 1X for 15 minutes. The coverslips were rinsed again three times in PBS and incubated with DAPI (1:1,000) in PBS for 10 minutes. The glass coverslips were rinsed in PBS three more time before being mounted on glass slides in Vectashield mounting medium. Binucleated cells with or without micronuclei were observed and counted under a fluorescence microscope. At least 500 binucleated cells were counted for each replicate condition. Statistical analysis was performed using Chi-squared tests with PRISM software.

### Statistics

All cell culture experiments were performed with at least three independent biological replicates unless otherwise indicated. RNA-seq experiments were conducted using four independent biological replicates, and RPPA analyses were performed using three independent biological replicates. Data are presented as mean ± standard deviation (SD) unless otherwise specified in the figure legends. Statistical analyses were performed using GraphPad Prism 10 (GraphPad Software) and R (version 4.x). For comparisons between two groups, unpaired two-tailed Student’s *t*-tests with or with Welch’s correction were used. Comparisons involving more than two groups were performed using one-way or two-way ANOVA followed by Tukey’s multiple-comparison test, or using Kruskal–Wallis tests followed by Dunn’s multiple-comparison test when data did not meet parametric assumptions. Differential transcription factor activity analyses from single-cell RNA-sequencing data were performed using Limma, while differential gene expression analyses of bulk RNA-sequencing data were performed using DESeq2 with Benjamini–Hochberg correction for multiple testing. Gene set enrichment analyses were conducted using fgsea, and adjusted *P* values were used to assess significance. Micronucleus frequencies were compared using Chi-square tests. All tests were two-sided unless otherwise stated. Differences were considered statistically significant when *P* < 0.05.

## Acknowledgements

We are grateful to DC Bennett and H Arnheiter for Melb-a cells and mi^vga9^ mice, respectively. Crispr/Cas9 constructs Thomas Zillinger (University Hospital Bonn, Germany). We acknowledge the animal colony (especially Pauline Dubreuil), histology (Sophie Leboucher), imaging (Marie-Noëlle Soler and Claire Lovo) and Christophe Roulin from RadexP facilities of Institut Curie.

## Authors contributions

J.H.R. conducted the bioinformatics studies, wrote the paper. M.L.C performed experiments and data analysis, and conducted the mouse studies. Z.A. carried out experimental work, analyzed the data, and performed the *in vivo* studies. M.H. conducted experiments and analysis. E.S. conducted experiments and analysis. V.F. conducted experiments and analysis. V.P. conducted experiments and analysis. A.B. interpreted the results. C.R.G. interpreted the results. L.L. designed the study, supervised the overall project, interpreted the results, wrote the paper.

## Ethics statement

All methods were performed in accordance with the relevant guidelines and regulations following good scientific practice.

Animal care and use for this study were performed in accordance with the recommendations of the European Community (2010/63/UE) for the care and use of laboratory animals. Experimental procedures were specifically approved by the ethics committee of the Institut Curie CEEA-IC#118 (Authorization #55477-2025040415461498 v2 given by National Authority) in compliance with international guidelines. Mice were housed in a specific-pathogen-free (SPF) certified animal facility with a 12-hour light/dark cycle in a temperature-and humidity-controlled room (22 ± 1 °C and 60%, respectively) with free access to water and food.

## Data availability statement

The RNA-sequencing data generated in this study have been deposited in the Gene Expression Omnibus. The publicly available datasets re-analysed here are available under accessions GSE236918 (mouse hair follicle) and GSE212450 (human scalp). All other data supporting the findings of this study are available from the corresponding author on reasonable request for non-commercial purposes.

